# Species coexistence and temporal environmental fluctuations: a quantitative comparison between stochastic and seasonal variations

**DOI:** 10.1101/2021.04.20.440706

**Authors:** Immanuel Meyer, Bnaya Steinmetz, Nadav M. Shnerb

## Abstract

Temporal environmental variations may promote diversity in communities of competing populations. Here we compare the effect of environmental stochasticity with the effect of periodic (e.g., seasonal) cycles, using analytic solutions and individual-based Monte-Carlo simulations. Even when stochasticity facilitates coexistence it still allows for rare sequences of bad years that may drive a population to extinction, therefore the stabilizing effect of periodic variations is stronger. Correspondingly, the mean time to extinction grows exponentially with community size in periodic environment and switch to power-law dependence under stochastic fluctuations. On the other hand, the number of temporal niches in periodic environment is typically lower, so as diversity increases stochastic temporal variations may support higher species richness.

## I. INTRODUCTION

Ecological and evolutionary processes take place in varying environments. As a result, demographic rates of populations fluctuate and their abundances vary through time [1]. An increase in demographic variability increases the chance of a population to reach the low-abundance region where extinction is plausible, so environmental variations have the potential to reduce species richness. One of the most surprising and counter-intuitive observations in community ecology has to do with the opposite effect: the ability of temporal environmental variations to *stabilize* a system of competing species and to *increase* the persistence time of diverse assemblages [2–6].

Diverse and even highly diverse communities of competing species are prevalent in nature [7–9], in apparent contradiction with the competitive exclusion principle [10, 11] and/or with the severe limitations exposed by May in his analysis of the complexity-diversity problem [12]. Therefore, the potential role of environmental variations in promoting taxonomic and genetic diversity received a lot of interest. Recently a few theoretical works provided analytical and numerical tools for a quantitative assessment of the stabilizing effect of environmental variations [5, 13, 14], and in parallel some prominent studies were focused on its manifestation in empirical communities [15–19]. The potential role of stochasticity-induced stabilization in protecting genetic polymorphism was considered as well [20–23].

An aspect that (as far as we know) did not receive an adequate attention is the distinction between periodic and stochastic variations. Although many natural environmental fluctuations are stochastic, most of them have at least some temporal periodicity or seasonality [24]. Quantities like yearly precipitation or the mean yearly temperature indeed fluctuate stochastically, but the amplitude of these stochastic variations is usually much smaller than the amplitude of the variations associated with the seasonal cycle during the year. Seasonal effects on demographic rates and on abundance and frequency variations are well-documented, both for populations [25] and for genotypes [26].

Despite a few exceptions [27–29], most of the studies of the relationships between temporal variability and stability were focused on stochastic variations. Here we would like to examine the periodic case and to contrast the outcomes with known results for stochastic environmental fluctuations. To do that, we will consider periodic and stochastic variations with the same characteristics (same amplitude of environmental fluctuations, same mean duration of environment’s dwell time, same level of demographic stochasticity) and will compare their stability properties. In two-species communities stability is quantified by the mean time to extinction, and in diverse communities with extinction-speciation equilibrium we implement species richness as a metric for the strength of the mechanisms that promote coexistence.

Following former works [30, 31], we consider individual-based version of the lottery model, which is the canonical example of Chesson’s storage effect [2–4]. To solve these models we implement the diffusion approximation, assuming that the characteristic timescales associated with environmental variations are much smaller that the time required for a given population to reach fixation [the “annealed” regime of Mustonen and Lässig [32]]. In the opposite (“quenched”) regime, the outcome of a competition is usually decided before an environmental shift occurs, so the role of the environmental variations in protecting coexistence is negligible.

Our analysis yields two main insights. First, all other things being equal, a community is more stable under periodic variations. Stochastic fluctuations, even when promoting coexistence, may drive a population to extinction through rare sequences of bad years, while seasonal cycles contribute only to stabilization. Second, in diverse communities the situation may change. Stabilization through temporal variations requires the number of temporal niches to be larger than the number of species, and seasonal cycles typically involve only a limited number of different states. Therefore, if stochastic variations offer much larger number of temporal niches they will play the main role in stabilizing diverse communities. The emerging picture and its relevance to practical implications are presented in the discussion section.

## II. TWO-SPECIES COMPETITION

Two-species competition is the elementary building block of coexistence theory. Here we implement a set of individual-based models, so our theoretical and numerical analyses take into account both demographic stochasticity (stochastic effects that influence the reproductive success of individuals in an uncorrelated manner) and environmental variability. All our models are simple continuous-time (Moran) generalizations of the classical lottery model (see [6] for details).

In an individual-based models global competition allow environmental variations to facilitate coexistence, while under dynamics with local competition those variations impede coexistence. Here we examine the differences between stochastic and periodic variations in both cases, so overall we consider four scenarios: global-periodic, global-stochastic, local-periodic and local-stochastic.

### A. Model definitions

We consider a zero-sum competition so the total number of individuals is *N*. The dynamic takes place in elementary birth-death events, and a generation is defined as *N* such elementary steps. Therefore, the duration of each elementary step is 1/*N*. The (time-dependent) fitness of a given population (species) is *ϵ^s^*, so *s* is the log-fitness or the selection parameter.

Due to environmental variations, *s* of each population is time dependent. In our models, the environment stays fixed for a certain time, that we define as its dwell time *δ*, and then it switches. *δ* is measured in generations, so if *δ* = 1, say, the typical dwell time of the environment is *N* elementary events.

- When the environmental variations are *periodic* or seasonal, the environment switches every *Nδ* elementary steps.
- When the environmental variations are *stochastic*, the chance of the environment to flip after each elementary step is 1/(*Nδ*), therefore, the dwell times are picked from an exponential distribution whose mean is δ generations.

Once the value of *s* for each individual is given, it determines its expected reproductive success. We study two competition scenarios.

- *Local competition* is perhaps the generic scenario for a community of animals who wander around in a certain spatial region, with an encounter between two individuals resulting in a struggle for a piece of food, territory and so on. To model such an interaction we consider dynamics that take place via a series of duels between two randomly-picked individuals, in which the loser dies and the winner reproduces. If the fitness of the first individual is exp(*s*_1_) and the fitness of the second is exp(*s*_2_), then the chance of the first to win the competition, *P*_1_, is determined by the fitness ratio

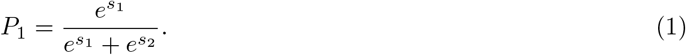 Analogously, *P*_2_ = 1 – *P*_1_.
- When the competition is global, one individual is chosen at random to die. The chance *P_a_* of an individual *a* to produce an offspring that recruits the resulting open gap is given by

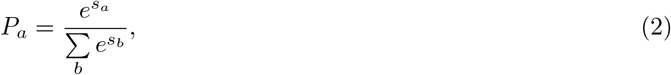

where the sum ranges over all the individuals. As a result, in the case when all the individuals in a population have the same fitness, the chance of species 1 (with *n*_1_ individuals) to capture the gap is

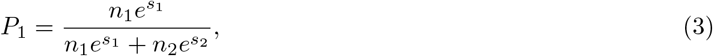

Since the fitness of each individual is either *s*_1_ or *s*_2_, Eqs. (1) and (3) imply that the outcome of the process depends only on the relative fitness Δ*s* = *s*_1_ – *s*_2_.

If *x* = *n*_1_/*N* is the frequency of species 1 (that we consider, without loss of generality, as the focal species), then the frequency of species 2 (rival species) is 1 – *x*. When species 1 wins, its frequency grows by 1/*N*. In the local competition model, the chance of an interspecific duel is 2*x*(1 – *x*), so the mean change in *x* after a single elementary event is

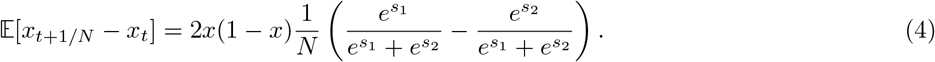

so when |Δ*s*| ≪ 1,

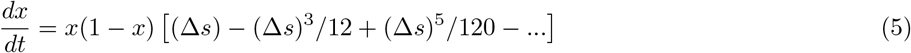

Importantly, the full expansion of the *dx/dt* in terms of Δ*s* admits only odd powers of Δ*s* multiplied by *x*(1 – *x*). When considering the logit parameter, *z* ≡ ln *x*/(1 – *x*), one notices that it satisfies *dz/dt* = (*dx/dt*)/[*x*(1 – *x*)]. Therefore, the changes in *z* when Δ*s* is positive are equal in magnitude and opposite in sign to the changes when Δ*s* is negative. As a result, under local competition fitness fluctuations cannot contribute to stability [33–35], they only lead to an unbiased random walk along the *z* axis.

The situation changes when competition is global. Now an increase (decrease) in the frequency *x* can happen if one of the rival (focal) species dies with probability 1 – *x* (*x*), and the slot is captured by the focal (rival) species. Therefore,

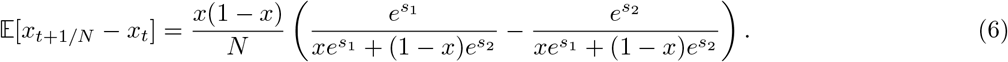

and,

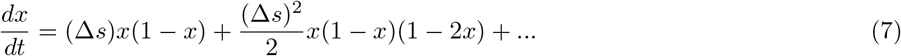

Under global competition, one finds in the expansion even terms, like (Δ*s*)^2^, that do not change sign when the relative fitness (Δ*s*) flips. The coefficient of (Δ*s*)^2^, the term *x*(1 – *x*)(1 – 2*x*), is positive at *x* < 1/2 and negative when *x* > 1/2, so it supports an attractive fixed point at *x* = 1/2. Therefore, under global competition variations in Δ*s* facilitate the invasion of rare species and stabilize the coexistence state, a phenomenon known as the storage effect.

To complete the picture, we have to quantify the effect of environmental variations, i.e., to specify how the selection parameter varies through time. In our models fitness variations are dichotomous, so (in case of two species) Δ*s* is either so + *γ* or *s*_0_ – *γ*. *s*_0_ is the mean fitness difference between the focal and the rival species (i.e., when *s*_0_ is positive the focal species is, on average, superior and vice versa), whereas *γ* is the amplitude of fitness variations. A glossary for all parameters is provided in Table I.

**TABLE I.**
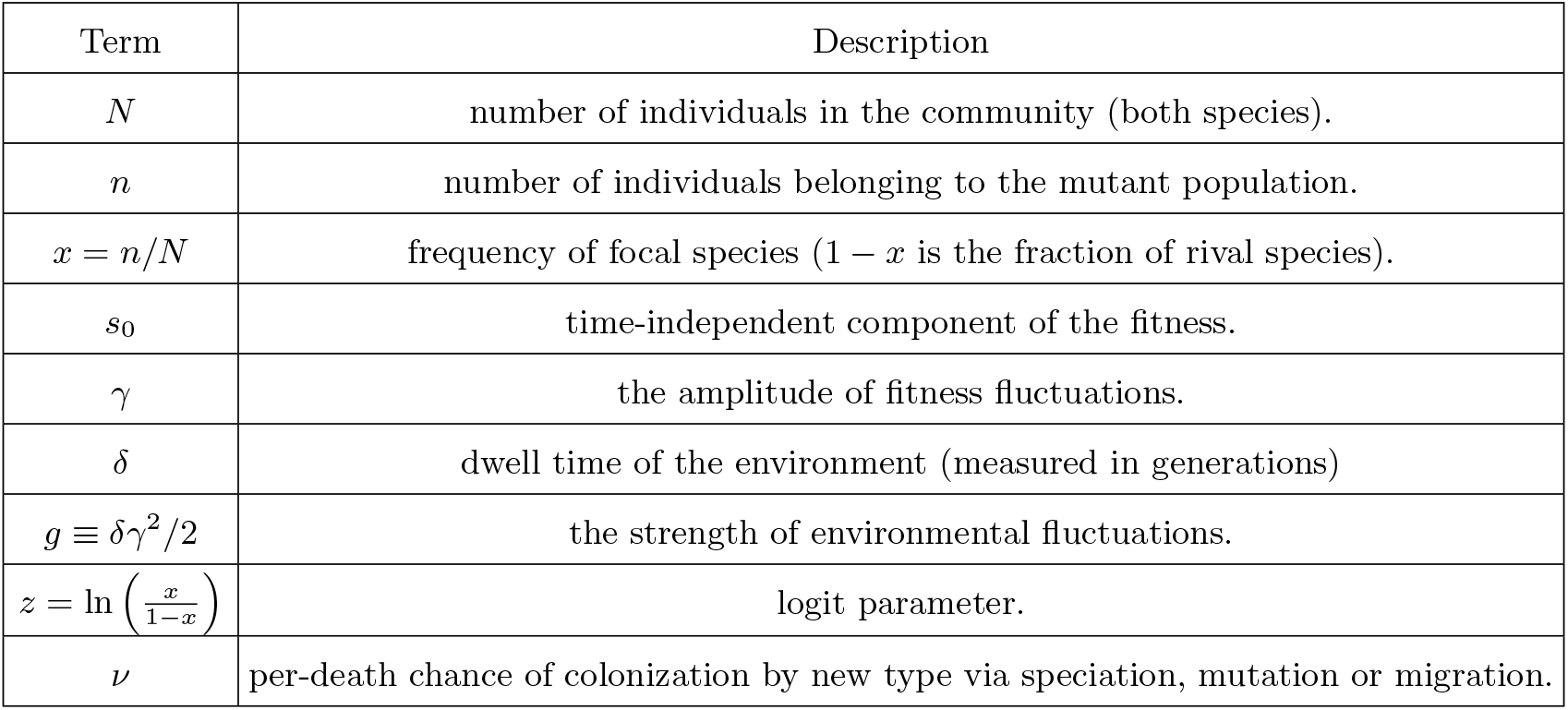
Glossary

### B. Analytic solutions

The main metric for persistence of a community is *T*, the mean time until one of the species is lost. We will refer to *T* as the time to extinction (of either species).

In our two-species community the state of the system at a given time is fully characterized by the frequency of the focal species *x* and the state of the environment. Since we consider only the annealed dynamics where *x* variations are much slower than environmental variations, we characterize the initial state of the community by *x* alone, so *T*(*x*) is the mean time until extinction, averaged over all initial states of the environment and over all histories [34].

Implementing the diffusion approximation one finds that *T* satisfies [34, 36, 37],

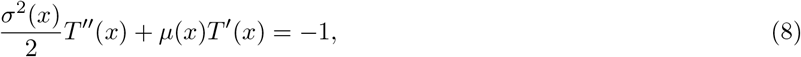

where *μ*(*x*) is the mean velocity, *σ*^2^(*x*) is the associated variance (see Methods, subsection V A) and prime represent a derivative with respect to *x*. Note that in Eq. (8) time is measured in elementary steps, to translate that into generations *T* must be divided by *N*.

In the Methods section (subsection V A) we calculate the values of *μ*(*x*) and *σ*^2^(*x*) for four combinations of global (*G*) and local (*L*) competition with stochastic (*S*) and periodic (*P*) environmental variations.

As shown in the previous section, when the competition is local the frequency preforms a random walk in the logit (*z*) space, where the right and the left moves are proportional to the fitness differences between the species. Therefore, when environmental variations are periodic the total displacement during a full cycle is independent of these variations, so *γ* and *δ* have no effect on *σ*^2^(*x*) and *μ*(*x*), which are simply those obtained in the standard case of a fixed environment,

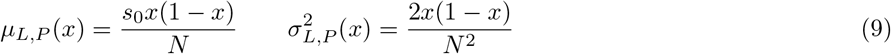

On the other hand, when the competition is global Eq. (7) suggests a bias towards the coexistence point. This bias is independent of the sign of Δ*s* and its strength is proportional to (Δ*s*)^2^, which when the diffusion approximation holds may be approximated by *γ*^2^. In the corresponding expression one finds an additional term in the expression for *μ*(*x*). This new term is proportional to *γ*^2^ and represents a bias towards *x* = 1/2.

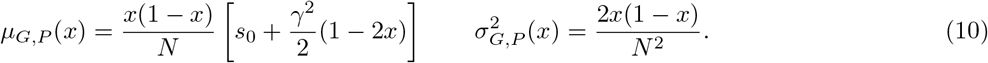

Since the variations are periodic, they do not contribute to the diffusion term *σ*(*x*).

When the environmental variations are stochastic abundance variability increases due to the chance of the focal species, say, to pick a sequence of good or bad years. In the local stochastic case the resulting parameters are,

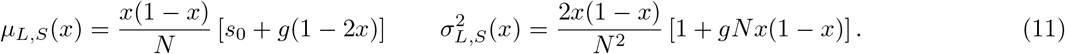

As before, the *μ*(*x*) has a term that support a stable fixed point at *x* = 1/2. However, the diffusivity *σ*^2^(*x*) is maximal at *x* = 1 /2 and vanishes close to the extinction point *x* = 0 and *x* = 1. The system tends to stick to the regions where its diffusion constant is small [6], and this “diffusive trapping” acts against the stabilizing effect of the *μ* term. These two contradicting tendencies cancel each other [6, 38], since in the logit space the dynamics is a simple random walk. The only net effect of stochastic environmental variations is an increase in the amplitude of abundance variations, which decreases extinction times [31, 34].

Finally, when competition is global the (Δ*s*)^2^ term of Eq. (7) breaks the tie in favor of stabilization and the storage effect manifests itself [34, 35],

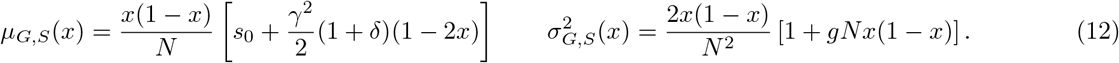

Eq. (8), with the relevant *μ*(*x*) and *σ*^2^(*x*), is an inhomogeneous and linear first order differential equation for *T*′, so one may solve it via an integration factor and then find *T* by an additional integration. When this procedure is implemented numerically, as detailed in the Methods section (subsection V B), the results fit perfectly the outcomes of our Monte-Carlo simulations, as demonstrated in Figure 1.

**FIG. 1.**
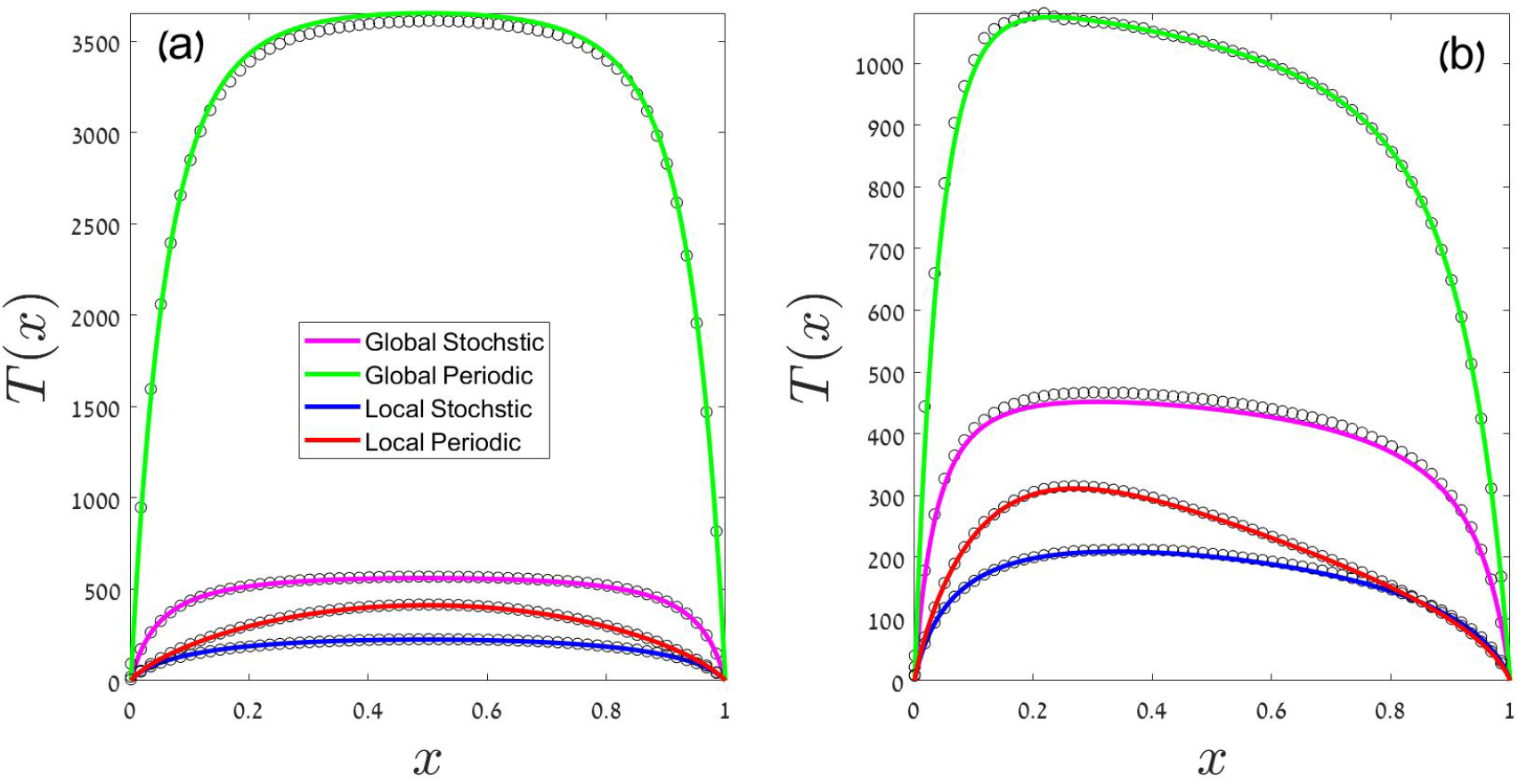
Mean time to extinction, *T*, is plotted against the initial frequency of the focal species *x*. Black circles are the outcomes of a Monte-Carlo simulation, and the colored lines are the theoretical predictions for the corresponding case (see legend) as obtained from numerical solutions of Eq. (8) with the *μ*(*x*) and *σ*^2^(*x*) from Eqs. (9-12). Details of the solution are presented in the Methods section. The hierarchy *T_GP_* > *T_GS_* > *T_LP_* > *T_LS_* is evident. In panel (a), both species have the same mean fitness (so = 0) and therefore all the lines are symmetric around *x* = 1/2. Other parameters in panel (a) are *N* = 600, 7 = 0.25 and *δ* = 0.55. In panel (b) the focal species is slightly advantageous, with so = 0.01, so the maximum time to extinction appears at *x* < 1/2. Other parameters in panel (b) are *N* = 500, 7 = 0.2 and *δ* = 0.1.

Figure 1 reveals the hierarchy of stability properties. The local-stochastic case has the shortest time to extinction, as the role of environmental variations is purely destabilizing. In the local-periodic case this destabilizing effect of environmental variations averages out over each cycle, so its *T* is larger. Under global competition temporal variations facilitate coexistence, so their *T* is larger than in the local cases. Finally, time to extinction in the global-stochastic scenario is shorter than in its global-periodic counterpart: a sequence of bad years may cause extinction in the former, but not in the latter case.

More quantitatively, the mean time to extinction in the local-stochastic and in the global-stochastic cases, and in addition *T* in fixed environment (which, as explained, is equivalent to the local-periodic case), were calculated by Danino *et al.* [34]. Let us quote some of their result, and contrast them with the new results obtained in the global-periodic case.

When *s*_0_ = 0, or otherwise in the weak selection regime where the effect of *s*_0_ is negligible, the maximum value of *T* (max over all values of initial state *x*) is,

- In the local-stochastic case, *T* ~ ln^2^ *N*.
- In the fixed environment or local-periodic case, the maximum value of *T* is linear in *N*. This is an old result, first obtained in Kimura [39]. Interestingly, Murray and Young [40] obtained this neutral-like results, for both the chace of ultimate fixation and the persistence time, using a model with *global* competition. They consider the parameter regime in which *γ*^2^ is negligible, so the expressions in (10) reduce to those of (9). Deviations from the neutral predictions are then observed only in the quenched regime of Mustonen and Lässig [32].
- In the global-stochastic case, *T* ~ *N*^1/*δ*^.

To understand these different dependencies on *N*, one notices that, as explained above, in the local-stochastic case with *s*_0_ =0 the abundance preforms an unbiased random walk along the logit (*z*) axis. By taking demographic stochasticity into account only as an absorbing boundary at single individual (*z* ≈ ±ln *N*), one finds exit times that scale with ln^2^ *N* [37]. When competition is global and the environment stochastic the *μ*(*x*) term support an attractive fixed point in *x* = 1/2, and extinction occurs through improbable sequences of bad years. During such a sequence abundance decreases exponentially, so the number of bad years required to cross the one individual threshold is proportional to ln *N*. Since the chance of such a sequence decreases exponentially with its length, the mean time to extinction scales like a power-law in *N*.

At last, in the global-periodic case (that was not discussed by Danino *et al.* [34]) there are no such rare sequences of bad years. Therefore, extinction may take place only due to demographic stochasticity (note that the *σ*^2^(*x*) term in Eq. (10) is the same as in Eq. (9), reflecting only demographic variations). This requires a highly improbable sequence of death events that may take a population with order *N* individuals to extinction, and such a sequence is exponentially rare in *N*. Therefore, one expects *T* in the global-periodic scenario to grow exponentially with *N*. These four behaviors: ln^2^ *N*, linear, power-law and exponential, are shown in Figure 2.

**FIG. 2.**
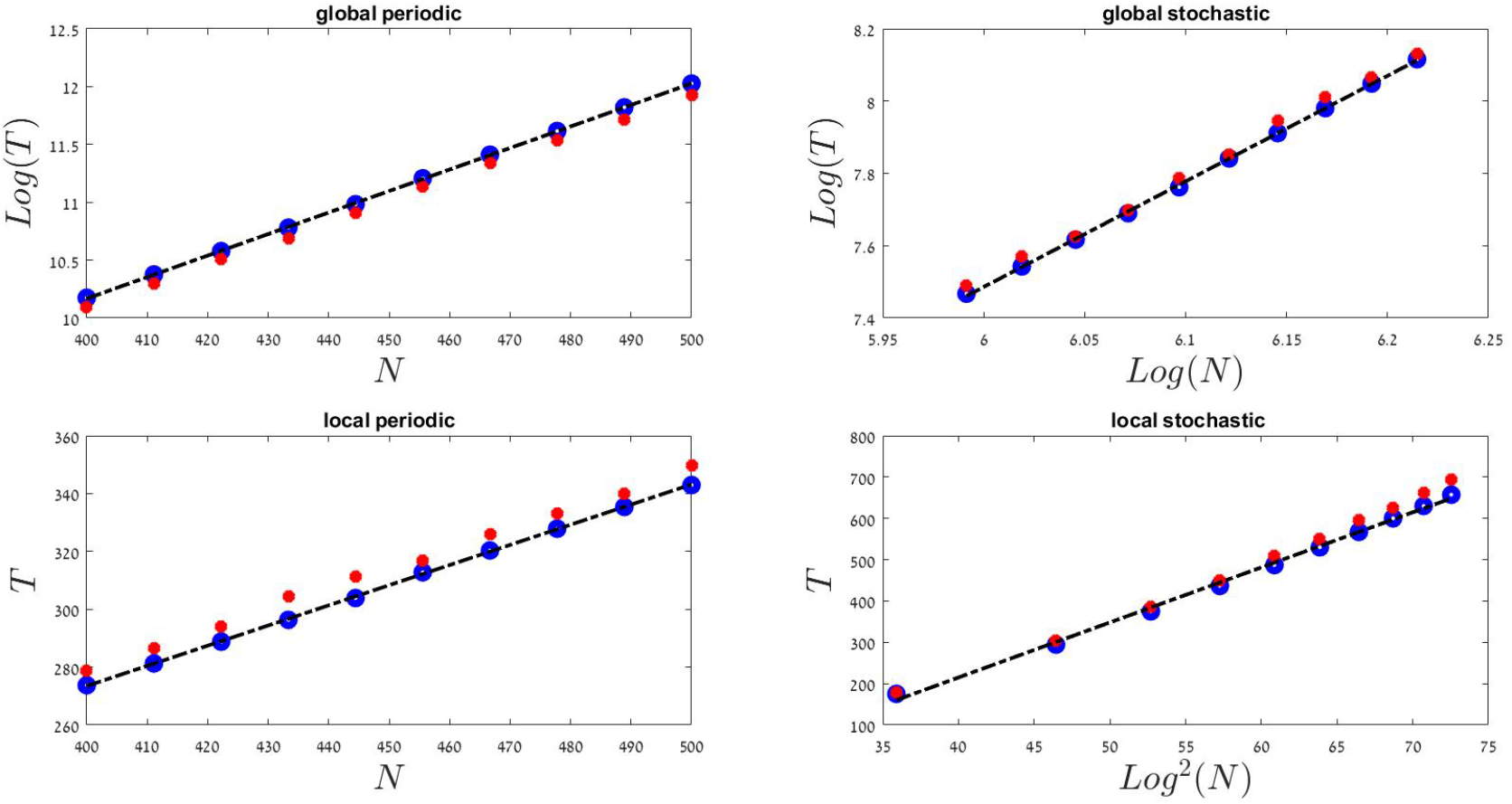
The relationships between the mean time to extinction *T* and the size of the community, *N*, in different scenarios. Filled red circles were obtained from Monte-Carlo simulations, Blue circles are theoretical predictions based on numerical integration of Eq. (8) with the *μ*(*x*) and *σ*^2^(*x*) from Eqs. (9-12), and the dashed black line is a linear fit to these blue circles, presented to guide the eye. As expected, in the global-periodic case the mean time to extinction grows exponentially with *N*, in the global-periodic the grows satisfies a power-law, the local-periodic case behaves like the neutral model (*T* is linear in *N*), whereas the local-stochastic dynamics yields log^2^ *N* growth. In all cases *s*_0_ =0 (so the *T* shown here is the mean time to extinction starting from *x* = 1/2), *δ* = 0.2 and 7 = 0.4.

When one cannot neglect so the situation is quantitatively different, but the qualitative hierarchy is preserved z[34]). In that case under local competition *T* ~ ln *N* (in both periodic and stochastic cases). When competition is global, as long as |*s*_0_| is not too large and *μ*(*x*) still posses an attractive fixed point at some 0 < *x* < 1, extinction takes place via accumulation of bad years (in the stochastic case) or random death events (in the periodic case) so *T* has the same scaling with *N*, although the actual time to extinction for a given set of *N*, *γ* and *δ* become shorter as |*s*_0_| grows.

### III. STOCHASTIC AND PERIODIC VARIATIONS IN DIVERSE COMMUNITIES

In the former sections we dealt with a two-species community. When environmental variations promote coexistence (global competition), seasonal cycles contribute only to the stabilizing mechanism. In contrast, when the fluctuations are stochastic a given population may pick a sequence of bad years that will take it to the brink of extinction. Therefore, all other things being equal, coexistence times under seasonal cycles were found to be longer than T under stochastic fluctuations.

In diverse communities this observation may change. Environmental variations facilitate coexistence only when different species in the community respond to the environment in distinctive manners, i.e., the stabilizing effect requires species-specific temporal niche. On the other hand, when a group of species has the same environmental response (fitness of different species changes coherently) the within-group relative fitness is kept fixed, so at least within this group there is no stabilizing effect.

Accordingly, the “species” that environmental variations promote their coexistence are not, or not necessarily, taxonomic species. More accurately, temporal fluctuations facilitate the coexistence of functional groups, i.e., of collections of organisms with similar traits that respond coherently to changes in temperature, precipitation and so on. Although the total abundance of each functional varies through time, variations in total abundance (like *N* variations) do not protect coexistence [41–44].

How many distinct temporal niches are allowed in a realistic system? The answer may reflect an essential difference between stochastic fluctuations and seasonal cycles. When the variations are stochastic a system may support, in principle, a large number of species, each with a specific temporal niche. Environmental stochasticity reflects erratic variations of a wide variety of factors that affect demographic rates - climate, predation pressure, resource availability, diseases and so on - and the different combinations of these factors suggest, potentially, many temporal niches.

Seasonality, in contrast, is usually associated with only a few environmental states. There is one profile of temperature and precipitation that characterises the summer, and another profile appears during the winter. Of course winters and summers properties varies among years, but these variations are not periodic; when winter-to-winter fluctuations are important, one has to include stochasticity in the model. Therefore, with respect to seasonal cycles one expects only a few temporal niches. Here we consider an extreme situation where only two temporal niches are available. These niches may stabilize two functional groups, species who are doing better during the summer and species that perform better during the winter. Environmental variations will stabilize the coexistence of these two groups, but will not promote coexistence within each group. Therefore, their overall effect on biodiversity turns out to be quite limited.

We stress that the comparison presented below, between the stabilizing effect of periodic cycles and stochastic temporal variations, is not a “fair” comparison. We compare systems with many temporal niches in the stochastic case, with systems with only two temporal niches in the periodic case. We would like demonstrate how periodic variations may lose their ability to promote coexistence when the number of temporal niches is small.

#### A. Models

In our simulations we implement the same set of competition models defined in Section II A for the two species case. We add to these models a new parameter, *v*, that reflects the rate in which new types (species, phenotypes) are added to the system via mutation, speciation or migration from a regional pool.

When an individual dies, in the two-species model the gap it leaves is recruited by an offspring of another individual, with probability that depends on fitness and other details of the dynamic. In the diverse community model this happens with probability 1 – *v*, and with probability *v* the gap is recruited by an originator of a new species.

Starting from a homogenous community (only one population) the number of species will increase until the rate in which new species are added to the system will be balanced by the rate of extinction. The equilibrium species richness *SR* serves as a metric of biodiversity. One expects *SR* to grow with the strength of the mechanisms that facilitate coexistence.

In stochastic environment, we assume that every new species is a new functional group, i.e., its response to environmental variations is uncorrelated with other species. Therefore after each environmental shift every species *i* picks its log-fitness *s_i_* at random and independently from a uniform distribution between –1/2 and 1/2. In periodic environments there are only two functional groups. When the environment shifts (transition from summer to winter and vice versa, *s_i_* → – *s_i_* for all the species in the system. Accordingly, species are divided into two general functional groups, those with higher *s* in summer and those with higher *s* in winter.

#### B. Results

As explained, all our simulations are initiated with a homogenous community with only one species. When the speciation rate *v* is very small, one has to wait for a long time until a new species appears. The global-periodic dynamic facilitates two-species coexistence better than the global-stochastic one. Accordingly, the invasion of the new species happens faster and the abundance variations are smaller in the periodic case. This behavior is demonstrated in Figure 3. As a result, when *vN* → 0 (which is analogous to the successive-fixation phase of [45], where one of the species goes extinct before the next speciation event happens) *SR* under periodic variations is larger, though its value is somewhere between one and two.

**FIG. 3.**
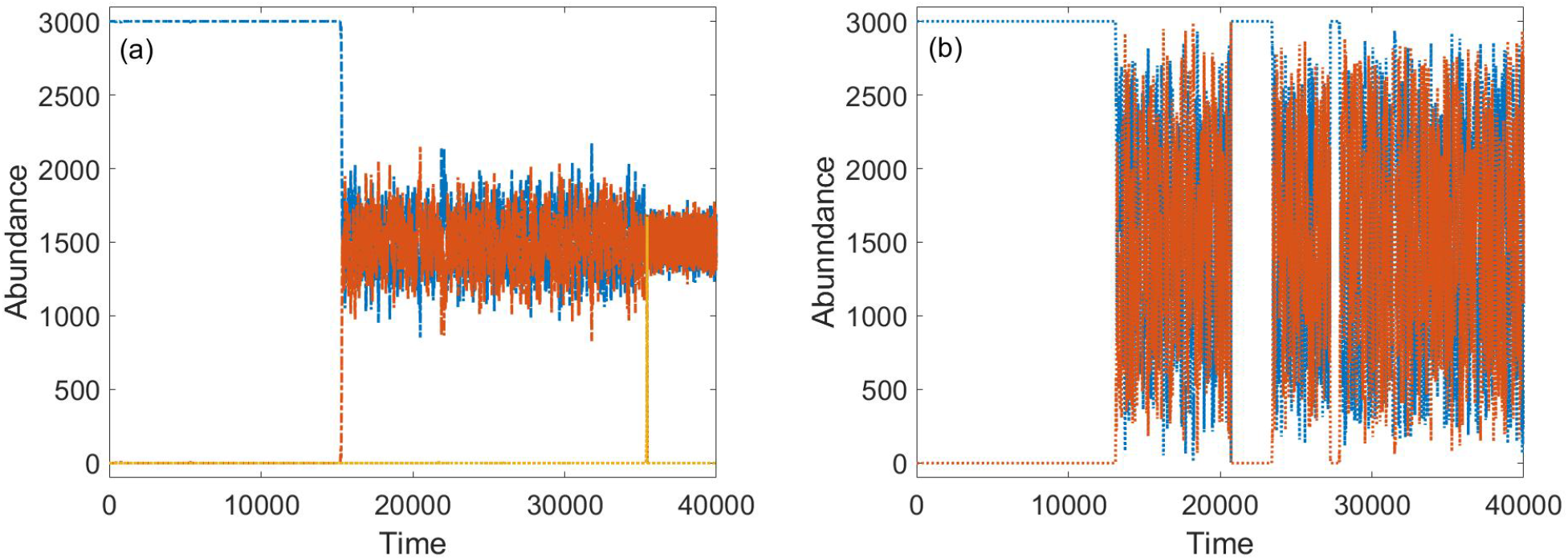
Community dynamics under global-periodic (panel a) and global-stochastic (panel b) competition. The abundance of different species in a community of *N* = 3000 individuals is shown for 40000 generations, where *δ* = 0.2 and *vN* = 0.001. The speciation rate is very small, so the resident species (blue) dominate the system for many generations before the first new species (red) invades. After invasion, periodic variations stabilize coexistence much better than stochastic fluctuations. In panel (b) abundance variations are larger, extinction events occur after long sequences of bad years and are followed by periods of monodominance. In contrast, in panel (a) the amplitude of abundance variations is much smaller and no extinction happens. Due to these stability properties, periodic systems are more diverse when *vN* is vanishingly small.

As the fundamental biodiversity parameter *vN* increases, the situation changes. Now many species try to invade. While in the stochastic case each species may find a time window in which it is fitter than all other species, in the seasonal system, as explained, there are only two type of populations and the fitness within each group is correlated. As exemplifies in Figure 4, as *vN* grows the richness of a stochastic system is higher than the richness of a periodic one.

**FIG. 4.**
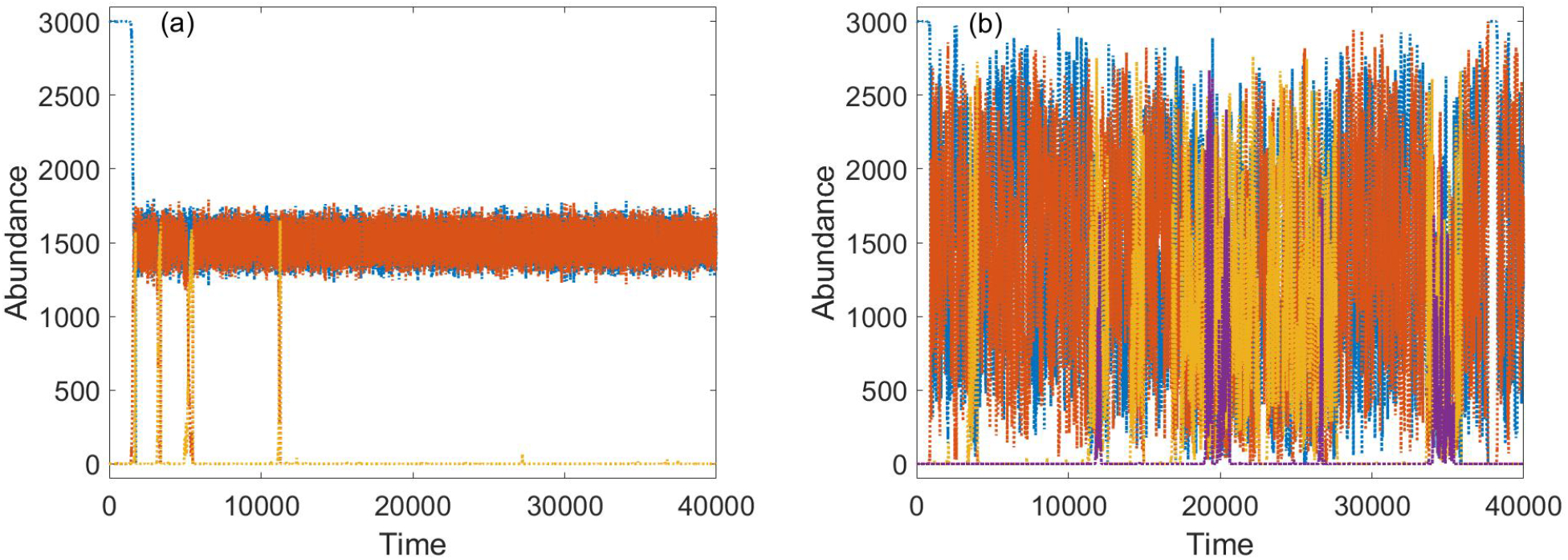
Same as Figure 3 except that *v* is larger by a factor of 10 so *vN* = 0.01. In the global periodic system (a), since there are only two temporal niches, every invasion attempt is readily followed by extinction of other species at the same functional group (note that colors are allocated to existing species only and when the red undergoes extinction, the yellow becomes red). In global-stochastic environment (panel b) abundance variations are larger, still more species coexist because the system admits an unlimited number of temporal niches.

The dependency of species richness on *N* when *vN* is large is summarized in Figure 5. Our baseline is the neutral model, where fitness variations vanishes and all species are demographically equivalent. Under local competition stochasticity destabilizes coexistence and diversity drops below the predictions of the neutral theory, whereas under global competition stochasticity promotes coexistence the species diversity increases (see [46] for a detailed comparison between stochastic systems). In contrast, in the periodic cases (both local and global) the species richness is more or less equal to the one obtained under neutral dynamics.

**FIG. 5.**
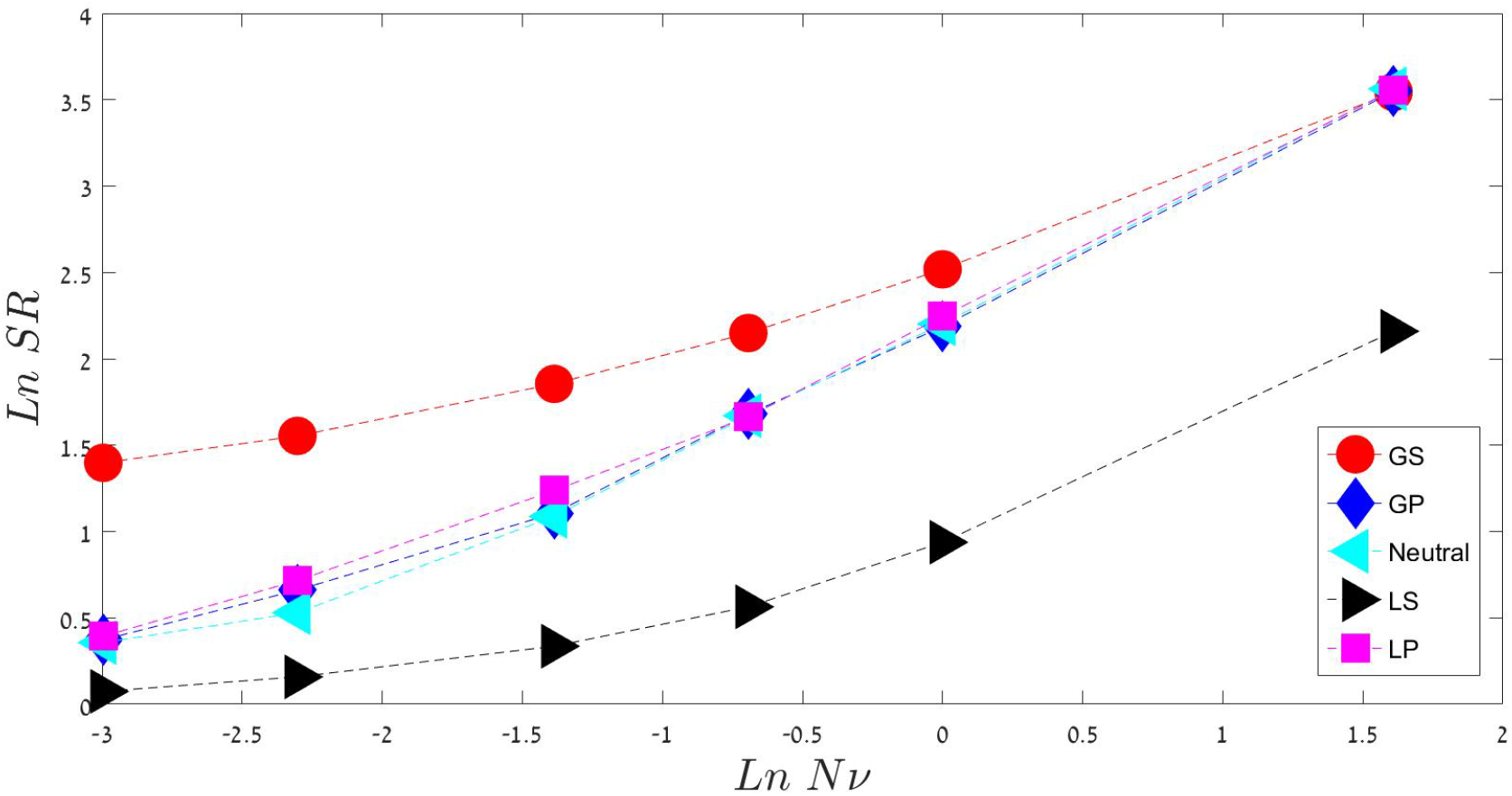
ln SR vs. ln *v*N in different scenarios of environmental variations. A community with global competition in stochastic environment, (GS), yields the highest species richness, while the cases of local competition in stochastic environment (LS), where temporal variations only impede coexistence, have the smallest diversity. Periodic variations with only two temporal niches, in either local (PL) or global (PG) competition modes, have essentially the species richness predicted by the neutral model, since within each functional group the dynamic is neutral. All results were obtained by averaging over long Monte-Carlo simulations with *N* = 5000 and *δ* = 0.2, where 7 was picked from a uniform distribution between −1/2 and 1/2. The fundamental biodiversity parameter *vN* ∈ [0.05..5].

To understand the convergence of the two periodic scenarios to the purely neutral limit, let us remaind ourselves that the species richness in a neutral system scales linearly with the size of the community *N* [47]. As explained, in a seasonal system species are divided into two functional groups, where the dynamics within each group is more or less neutral. Therefore, the whole community is made of two competing functional groups with neutral dynamic. Within each group *SR* is linear in its abundance, so overall the total richness will be the same. The actual abundance of each functional group fluctuates around *N*/2, but the effect of these fluctuations (unless they are extremely large and lead to bottlenecks that reduces diversity) is quite marginal.

## IV. DISCUSSION

Seasonality, and the temporal population variability associated with it, were observed in a wide variety of empirical systems [24–26]. Through this paper we provided a survey of its potential role in the maintenance of biodiversity in ecological communities.

We showed that periodic variations promote coexistence better than stochastic fluctuations, since in a periodic environment one cannot find rare sequences of bad years that drive populations to extinction. More quantitatively, whereas facilitation of two-species coexistence state by stochastic abundance fluctuations yields persistence times that grow like a power-law in *N*, under periodic variations yields *T* grows exponentially with *N*.

These results were demonstrated only for a two-species community, but one expects it to hold even when the number of species is larger, provided that each species admits its own temporal niche. In diverse communities this requires a set of narrow width niches that preserve a rigid temporal order. As species richness grows, these conditions become less and less plausible.

To examine the case where the number of temporal niches is smaller that the number of species, we consider the richness at an extinction-speciation equilibrium. Stochastic fluctuations (where the number of temporal niches is assumed to be unlimited) are compared to seasonal variations with only two distinct temporal niches (summer and winter, say), where the baseline is the equilibrium richness of Hubbell-Kimura neutral model [47, 48].

We discovered that a system with only two niches may stabilize only two functional groups, where within each group the dynamics is more or less neutral. Accordingly, when the typical richness of the system (roughly measured by the fundamental biodiversity parameter *vN*) is larger than the number of temporal niches, the diversity of the community sticks to its neutral value. Again, although we have examined numerically this case only in a community with two temporal niches, the main insights seem to be generic.

As always, typical natural scenarios are likely to be more complicated. For example, most systems are affected by both stochastic and periodic variations, although the amplitude of periodic variations (in temperature or precipitation between summer and winter, say) is usually much larger that the year to year stochastic fluctuations. The number of temporal niches under seasonal forcing may be larger than two: phenological differences may allow for a wide variety of temperature niches, as described by Mellard *et al.* [25]. Still, we believe that the results presented in this work provide the essential insights required in assessing the relative importance of these effects and their potential contribution to the richness of life forms in our world.

## V. METHODS

### A. Derivation of *μ*(*x*) and *σ*(*x*) in various cases - the diffusion approximation

As explained in the main text, in case of local competition, the deterministic change in *x* per unit time of a single step 1/*N* (its instantaneous velocity) is,

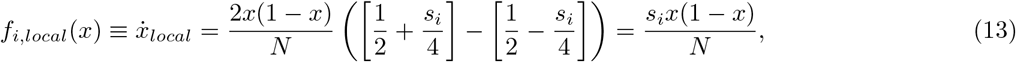

while for global competition,

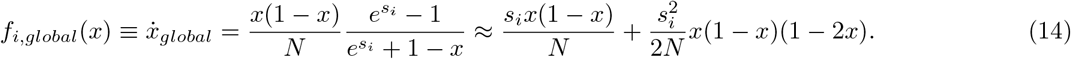

In addition, every single birth-death event is associated with a variance (demographic fluctuation) of 2*x*(1 – *x*).

In both cases, after *k* steps the mean change in *x* (for a given history of the selection parameter *s*(*t*)) will be,

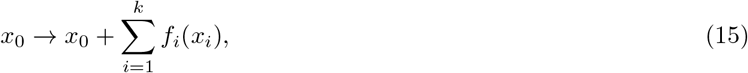

where *f_i_* is the velocity in a given *i* environmental situation (global or local). Expanding this sum to first order around *x*_0_, one gets,

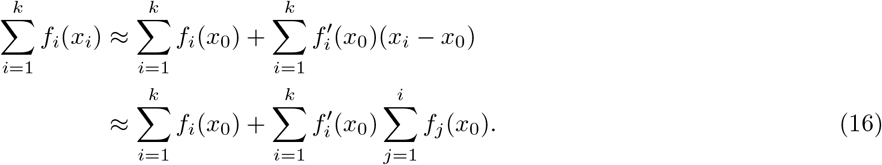

When the environmental fluctuations are periodic, **every** positive period is followed by a negative one of the same amplitude and duration and the internal summation is reduced to order *s*_0_. Once multiplied by the external summation, this term is negligible. Therefore, the mean displacement over several periods is,

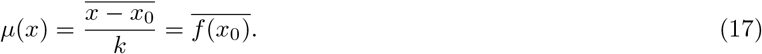

Plugging in (13) and (14) yields,

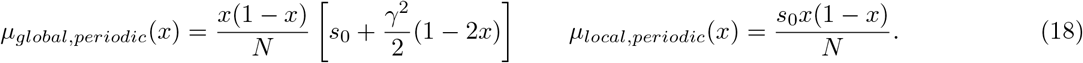

The square of the displacement is,

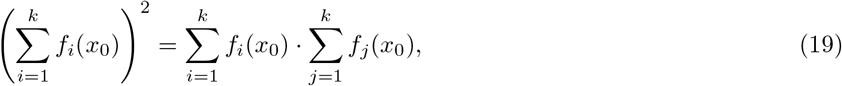

and this quantity vanishes, as before, over an integer number of periods. Therefore, the only contribution to *σ*^2^(*x*) comes from demographic stochasticity:

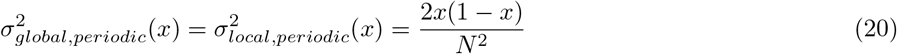

For the stochastic cases we can use the assumption that the environmental states are uncorrelated over periods that are longer than the dwell time, and simplify (16) by grouping together elementary steps into periods in which the environment is fixed,

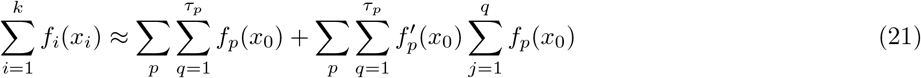

In this expression, the environment is stable during periods of *τ_p_*, and we sum over all environmental states indexed *p* during the *k*-th step. Since the *f_p_* of different *p* are uncorrelated, the internal summation of Eq. (16) is reduced to the sum over steps of the same state. Therefore,

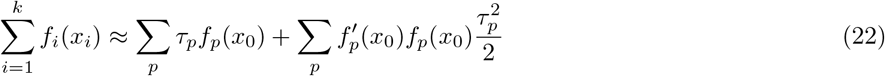

When the values of *τ_p_* are drawn from an exponential distribution whose mean is *τ*, 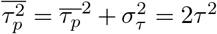, where 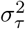 is the variance of *τ_p_*. Using,

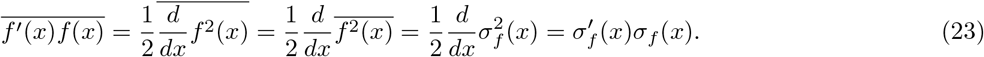

since in the local case, and approximately in the global case,

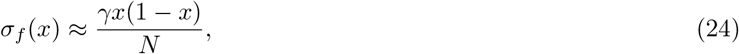

one can write

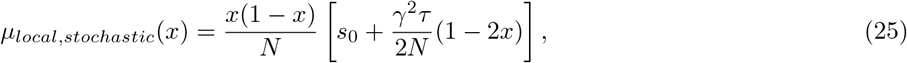

and

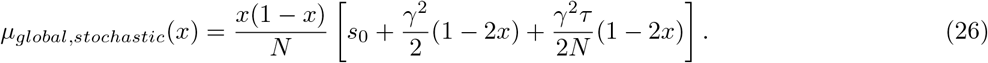

Finally, the variance in the stochastic cases is:

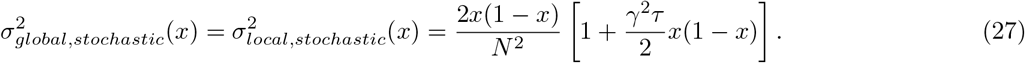

At this point we define *δ*, which is the environmental correlation time measured in generation. If a generation is defined as *N* elementary birth-death events, and on average every *τ* elementary birth-death steps the environment changes, then *δ*=*τ/N*. Correspondingly we define *g* = *γ*^2^*δ*/2 as the strength of the environmental variations.

### B. Solutions for *T* using integration factor

Eq. (8) of the main text reads,

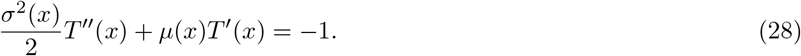

We solve it by using an integrating factor, such that

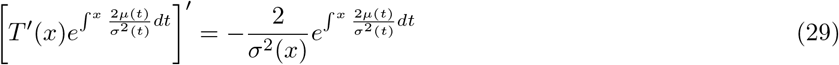

Integrating on both sides yields

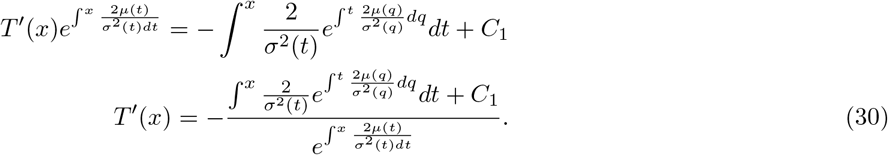

Finally,

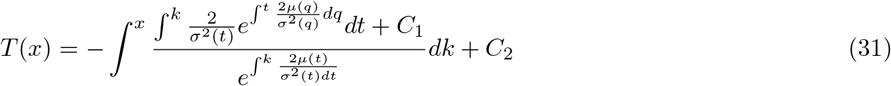

All that is left is to find the constants using the boundary conditions *T*(0) = *T*(1) = 0.

## VI. ACKNOWLEDGMENTS

This research was supported by the ISF-NRF Singapore joint research program (grant number 2669/17).The work of I.M. was supported by an Eshkol Fellowship of the Israeli Ministry of Science.

